# Future COVID19 surges prediction based on SARS-CoV-2 mutations surveillance

**DOI:** 10.1101/2022.09.05.506640

**Authors:** Fares Z. Najar, Evan Linde, Chelsea Murphy, Veniamin A. Borin, Huan Wang, Shozeb Haider, Pratul K. Agarwal

**Affiliations:** High-Performance Computing Center, Oklahoma State University, Stillwater, Oklahoma; Department of Physiological Sciences, Oklahoma State University, Stillwater, Oklahoma; University College London School of Pharmacy, Pharmaceutical and Biological Chemistry, London, United Kingdom; University College London Centre for Advanced Research Computing, London, United Kingdom

## Abstract

COVID19 has aptly revealed that airborne viruses such as SARS-CoV-2 with the ability to rapidly mutate, combined with high rates of transmission and fatality can cause a deadly world-wide pandemic in a matter of weeks.^1^ Apart from vaccines and post-infection treatment options, strategies for preparedness will be vital in responding to the current and future pandemics. Therefore, there is wide interest in approaches that allow predictions of increase in infections (“surges”) before they occur. We describe here real time genomic surveillance particularly based on mutation analysis, of viral proteins as a methodology for *a priori* determination of surge in number of infection cases. The full results are available for SARS-CoV-2 at http://pandemics.okstate.edu/covid19/, and are updated daily as new virus sequences become available. This approach is generic and will also be applicable to other pathogens.

## INTRODUCTION

Protein and genome sequence analyses identify molecular level changes that enable viral adaptations for increased spread through the host population. Concrete evidence for direct a relationship between specific mutations and increase in rates of infection (and fatality) requires extensive laboratory studies that need significant time. The availability of unprecedented number of SARS-CoV-2 genome sequences is making possible identification of number and types of mutations, which in turn can provide vital knowledge in real time, crucial for decision making by health professionals for medical interventions. We are investigating several different approaches (synonymous, non-synonymous, or non-synonymous/ synonymous ratio for the nucleotide sequences,^2^ and the conservation or radical substitutions for the amino acid sequences) for using number and types of mutations as a means to predict surge in infections as well as monitor the changes in critical viral proteins. Recently, such analysis has been reported for single SARS-CoV-2 proteins.^3^ Our approach, however, is based on analysis at the whole viral genome level and moreover it is performed continually in real time.

## RESULTS

It was found that collective non-synonymous mutations in key proteins of SARS-CoV-2 showed significant increase 10-14 days before the rapid rise in COVID19 cases, particularly related to the surges that occurred after the emergence of Gamma, Delta, Omicron and BA5 variants (Figure 1). At present, over 6.1 million SARS-CoV-2 genome sequences collected all over the world are available from GenBank (https://www.ncbi.nlm.nih.gov/sars-cov-2/), which were used for analysis of 26 SARS-CoV-2 proteins including the structural (spike, envelope, membrane, nucleocapsid) proteins, non-structural proteins (NSPs) and open reading frames (ORFs). Note, our analysis was performed with the first reported (“Wuhan”) SARS-CoV-2 sequence as a reference.^4^ In other words, the computed mutations are calculated in comparison to this reference sequence. The reason for an increase in mutations ahead of a surge is the search for adaptation against the acquired immunity (or gain in function) in either a single protein or a combination of proteins. The case of the Omicron variant indicates the development of the most drastic changes in several different proteins, which coincided with the largest increase in rate of infections (Figure 1). Non-synonymous mutations (k_a_) in several proteins show significant increase before the increase in rate of infections (or surges), therefore, allowing a means for surge prediction. Note that the ratio of non-synonymous to synonymous mutations (k_a_/k_s_) which is typically used, did not provide clear trends; furthermore, the rate of mutations (derivative of observed mutations wrt time) also did not a provide reliable prediction signal (see the supporting information for details). Below we provide a summary of the key results and their importance.

**Figure 1:**
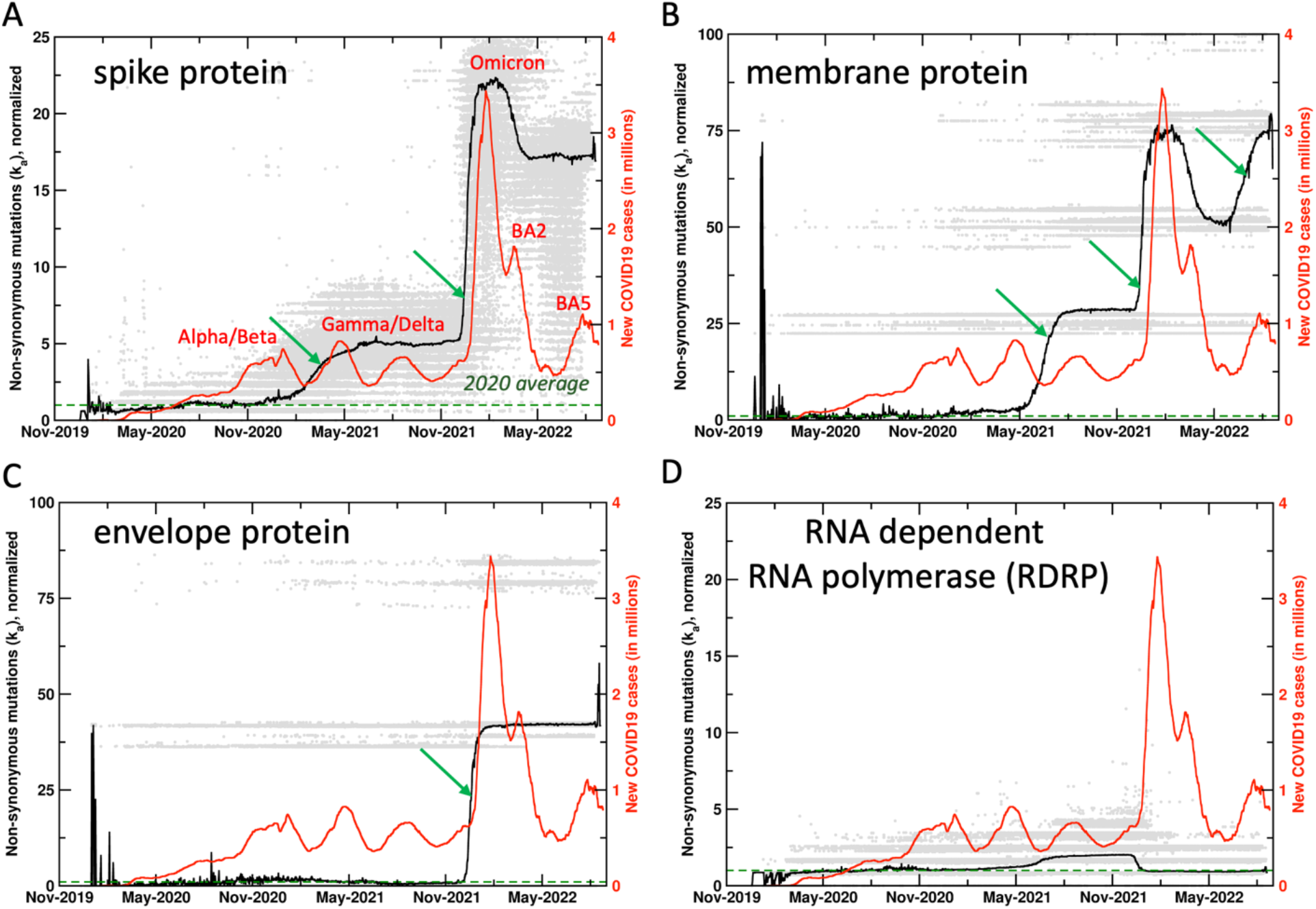
Mutations in SARS-CoV-2 proteins increase before COVID19 surges. Non-synonymous mutations over the course of the COVID19 outbreak were identified by analysis of 5.9 million sequences. Gray dots indicate individual mutations, while black lines show weighted means for each day. Red lines show new COVID19 cases (averaged weekly) across the world. The green arrows mark the time when new mutations occurred before the outbreaks, allowing prediction of future outbreaks. The mutation values have been normalized using average of all mutations in the year 2020 (the first full year of the pandemic) as 1 (marked by dashed lines). Raw results are available in supporting information. Values of 0 indicate same sequence as the Wuhan sequence, while larger values indicate more mutations. Note that each gray dot corresponds to a unique sequence, and there can be multiple sequences showing the same mutations. The weighted mean for the day is calculated by using all sequences reported for the day. The peaks for COVID19 cases are labeled with prevalent variants. Alpha/Beta, Omicron and Omicron BA2, BA5 were the prevalent variants at the time of labeled peaks. For the two peaks in 2021 the case was less clear, with Gamma and Delta variants being observed at different times in different parts of the world.

### Spike Protein

Spike protein interacts with the angiotensin-converting enzyme 2 receptor and plays a vital role in infecting the human cells.^5^ Spike protein has been the target of mRNA-based vaccines. Viral sequences show significant changes in synonymous and non-synonymous mutations in the spike protein (169,892 unique sequences observed so far), with large increases ahead of the surge in reported human infections, most noticeably with the surges associated with the Gamma/Delta and the Omicron variants (Figure 1A). It is important to note that the mutations show increase 10-14 days before the increase in human infections. It is also interesting to note that the synonymous mutations (data available on the website) show decrease post surges. The decrease in mutations prior to the Omicron BA2 surge corresponds to reversal mutations (returning to reference sequence). However, at present the non-synonymous and synonymous mutations post the Omicron variant remain elevated, more so than any period during the COVID19 outbreak.

### Proteins showing significant mutations

In addition to the spike protein, SARS-CoV-2 membrane^6^ (Figure 1B, 10,285 unique sequences observed so far) and envelope^7^ (Figure 1C, 1,221 unique sequences observed so far) proteins have also shown significant mutations, starting just before the Omicron variant (November 2021 onwards). For the case of membrane protein, there was significant increase that started in the Gamma/Delta variants (June 2021 onwards) and further increased just before the BA5 surge. The spike, membrane and envelope proteins are all located on the surface of SARS-CoV-2 and potentially interact with the components of the immune system. The large increase in mutations in all these external proteins assumes importance in post-vaccination period (discussed further below).

### Other proteins

For comparison, Figure 1D shows mutations from RNA-dependent RNA polymerase (RDRP, 55,531 unique sequences observed so far), which has been targeted for development of antiviral drug therapies. Till present, RDRP has shown comparatively lower magnitude of non-synonymous mutations. Note that gray dots are individual mutations, the mean (black line) is weighted by number of sequences for each day by the mutations. Significant increases in mutations are also observed in NSPs 1, 4, 6, 13, 15, ORFs 6, 7a and 7b (data available on website). Overall, this analysis allows us to monitor ongoing mutations in different proteins; when rapid rise is observed over a short period of time, we issue surveillance warnings for new possible variants (with combination of proteins showing new mutations).

### Vaccination and mutational frequencies

Wide-spread vaccination against SARS-CoV-2 (December 2020 onwards) coincides with significant increase in mutation rates of several SARS-CoV-2 proteins. Spike, membrane and envelope proteins have shown rapid mutations in especially in the Omicron variant (gray dots in Figure 1). This is possibly due to viral adaptations under the selective pressure exerted by the vaccine, as a significant number of mutations were observed in 2021, especially for the spike protein (gray dots in Figure 1A indicate spike protein has mutated much more than any other protein). The long-term effectiveness of mRNA-based SARS-CoV-2 vaccines remains unknown. After the initial regimen of two doses, the administration of additional booster (third and fourth) doses has decreased due to improvement in COVID19 fatality rates as well as political reasons. This situation raises concerns. Other proteins have shown reversal mutations (higher similarity with the reference sequence) after periods of significant increase in mutations, however, post-vaccination the significant mutations observed in the spike, envelope and membrane protein related to the Omicron variant remain at extremely elevated levels. As Omicron, BA2 and BA5 variants are showing increased rates of transmission, gain or improvement of function in other proteins could lead to emergence of newer variants of concern. Over long-term this needs to be addressed by vaccines with longer periods of effectiveness and post-infection treatment options including antiviral drugs.

### Surge prediction

The methodology presented here allows monitoring the potential increase in reported number of human infections. To date, spike protein has shown the most direct correlation in the rate of non-synonymous mutations and the rates of human infection. In particular, the case of Omicron variant and also the Gamma variant, spike protein showed rapid increase in mutations about 10-14 days ahead of time. Furthermore, membrane protein showed rapid mutations before surge related to BA5. Therefore, such increase is mutations serve as an indication of upcoming surges. For example, we issued a surge watch on the website on June 29^th^ 2022, which was confirmed in July.

The role of different (or dominant) SARS-CoV-2 variants in major surges is unclear at this time and needs further research. Different variants have been prevalent in different geographic regions at different times over the course of COVID19 outbreak, therefore, it is difficult to assign the surges to individual variants. In particular, Gamma and Delta variants were both prevalent in different countries in 2021. We are working on enabling this analysis by geographic locations and the results will be available through the website. However, at present our analysis is able to make predictions about collective surges before they occur, as illustrated by the case of BA5.

## SUMMARY

The methodology and the website described here provides real time mutational changes of 26 SARS-CoV-2 proteins and ORFs. The changes in non-synonymous mutations correlate with the increase in reported cases of infections. Apart from identifying mutations of concerns for in-depth scientific studies, the website is intended to keep the medical community informed about the potential surges. Warnings of increase in mutations and expected surges are displayed on the website (and also available through email alerts). It should be noted that this real time analysis is dependent on the various health labs and medical facilities for swiftly depositing the viral genome sequences into the public databases such as the GenBank. The shorter the lag time in depositing the sequences by the wider community, more accurate and effective the prediction capabilities of our approach and the website will be.

