## Supporting Information for "Future COVID19 surges prediction based on SARS-CoV-2 mutations surveillance"

### Methods

The SARS-CoV-2 genomic sequences data and the number of COVID19 sequences are continually obtained from the sources described below. The genomic sequences are carefully filtered for quality control and used for calculations of non-synonymous ($k_a$) and synonymous ($k_s$) mutation rates for each of the listed 26 proteins separately.

*Data and data sources:* Data for the number of reported COVID19 cases was accessed from Johns Hopkins University's Our World In Data project (https://ourworldindata.org/coronavirus-source-data).[1]

*Genomic sequence data:* An in-house pipeline of scripts (using Linux commands) was designed around the eUtils tools[2] from NCBI in order to download and process the SARS-CoV-2 records from NCBI's GenBank (https://www.ncbi.nlm.nih.gov/genbank/). Briefly, we used `esearch` and `efetch` commands to obtain these GenBank records. Search string "SARS-CoV-2", refined to "SARS-CoV-2 [ORGN]", was used to download the identified records in the GenBank text format. After workflow optimization, post May 2022, the search process used NCBI's newer `datasets` and `dataformat` command-line tools to identify sequences of interest while continuing to use the `efetch` tool to download records in the GenBank text format. Collectively, a total of 6,120,032 records were searched and a total of 2,865,328 sequences matching the search criterion were downloaded and used as of August 23rd, 2022.

*Quality control*: Incomplete and ambiguous SARS-CoV-2 genomic sequences and records containing incomplete collection dates were filtered out using the designed pipeline. For the records passing the quality control steps, the nucleotide sequence for each gene was extracted. A non-redundant version of the extracted nucleotide sequences was derived and translated to the cognate amino acids sequences. In the final phase of the pipeline, the accession numbers for each viral isolate with the nucleotide sequences, the associated protein sequences, the collection dates, and the country of collection were stored in SQLite relational database where they were indexed with unique identifiers to allow the retrieval and analysis of any part of the parsed data.

*Frequency of data updates:* As of July 2022, the described sources are monitored daily for updates. New data is continually downloaded and used for analysis.

*Alignments and non-synonymous ($k_a$), synonymous ($k_s$) calculations*: The translated proteins and nucleotides sequences were aligned using clustal-omega[3] and Pal2Nal[4] programs to align the codons with their associated amino acids. The resulting alignments were then processed through the program kaks_calculator[5] to calculate and non-synonymous ($k_a$), and synonymous ($k_s$), and their ratio $k_a/k_s$ values which were used to assess the mutational adaptation for each protein. The parameters required for the kaks_calculator were based on the maximum-likelihood method derived from the work of Goldman and Yang.[6] The first reported SARS-CoV-2 genomic sequence ("the Wuhan sequence")[7] was used as a reference for all the $k_a$, $k_s$ and $k_a/k_s$ calculations. We explored the possibility of using other sequence(s) as references (for example, the previous day or the previous month), however, due to the increasing number of variations available every day, it is difficult to select a representative sequence on an ongoing basis. It was also found that using the Wuhan sequence as a reference provided the most intuitive and interpretable results.

*Use of mutational rates as a surge predictor*: In addition to using collective number of mutations (Figure 1 in main text and Figure S1), commonly used $k_a/k_s$ ratio (Figure S2) and the rate of mutations (Figure 3) was also explored for suitability of surge prediction. As shown in Figure 3, $k_a/k_s$ did not provide a reliance surge prediction signal. Figure S3 shows rate of mutations (calculated as a numerical derivative). For the case of Omicron surge, the proteins did show increased rate of mutations, however, for all other cases a clear signal was absent. Furthermore, the rate of mutations approach presented two additional challenges. First, a number of instances were noted where the rate of mutations increased that did not precede surge in reported infections. Second, the most recent incoming genomic data is noisy (due to smaller number of samples and weighting of different mutations shows large variations) and changes quickly, therefore, the ongoing most recent rate of mutations is very noisy as well. It was concluded that at this stage, rate (derivative) of mutations is not a reliable signal for surge prediction. In the future, this could be revisited with more stable reporting of genomic sequences with shorter sample collection to sequence publication timeframes.

**List of proteins investigated**

The number of unique nucleotide sequences observed till date for each of the 26 proteins/open reading frames are listed below. The full results are available on the project website https://pandemics.okstate.edu/covid19/ Only three proteins showing the most relevant results and one protein (marked by *) for comparison is depicted in Figures. These proteins are shown in bold below.

1. **Envelope protein: 1,221**
2. **Membrane protein: 10,285**
3. Nucleocapsid protein: 64,459
4. **Spike protein: 169,892**
5. Non-structural protein 1 (NSP1), leader protein: 10,188
6. NSP2: 61,822
7. NSP3: 221,696
8. NSP4: 29,071
9. NSP5, 3C-like Proteinase: 11,066
10. NSP6: 15,572
11. NSP7: 1,237
12. NSP8: 3,933
13. NSP9: 2,701
14. NSP10: 2,272
15. NSP11: 87
16. **NSP12, RNA-dependent RNA polymerase (RDRP)*: 55,531**
17. NSP13, helicase: 31,755
18. NSP14, 3'-to-5' exonuclease: 26,124
19. NSP15, endoRNAse: 12,196

20. NSP16, 2'-O-ribose methyltransferase: 7,197
21. ORF3a: 39,216
22. ORF6: 2,015
23. ORF7a: 8,408
24. ORF7b: 1,295
25. ORF8: 6,582
26. ORF10: 670

Total number of quality-controlled SARS-CoV-2 sequences analyzed: 2,731,572 (as of August 23rd, 2022).

### Raw results (without normalization)

The un-normalized non-synonymous mutations ($k_a$) results are shown in the Figure S1.

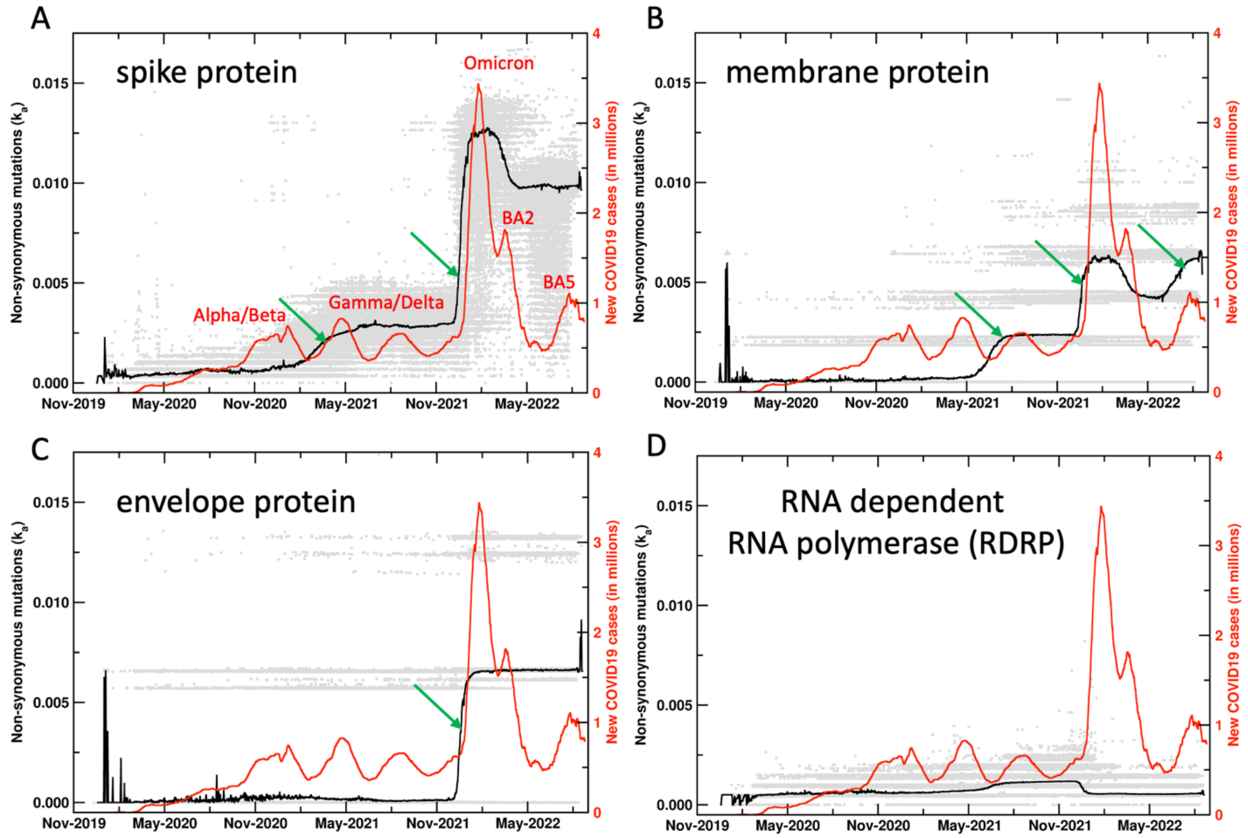


**Figure S1: Un-normalized results for the mutations in SARS-CoV-2 proteins.** See Figure 1 in the main manuscript for more details. Here the raw results for the four proteins are plotted for the non-synonymous mutations. The same y-axis scale is used for comparison of the mutations across all the four proteins shown.

**$k_a/k_s$ results**

The results for commonly used ratio $k_a/k_s$ are shown in the Figure S2.

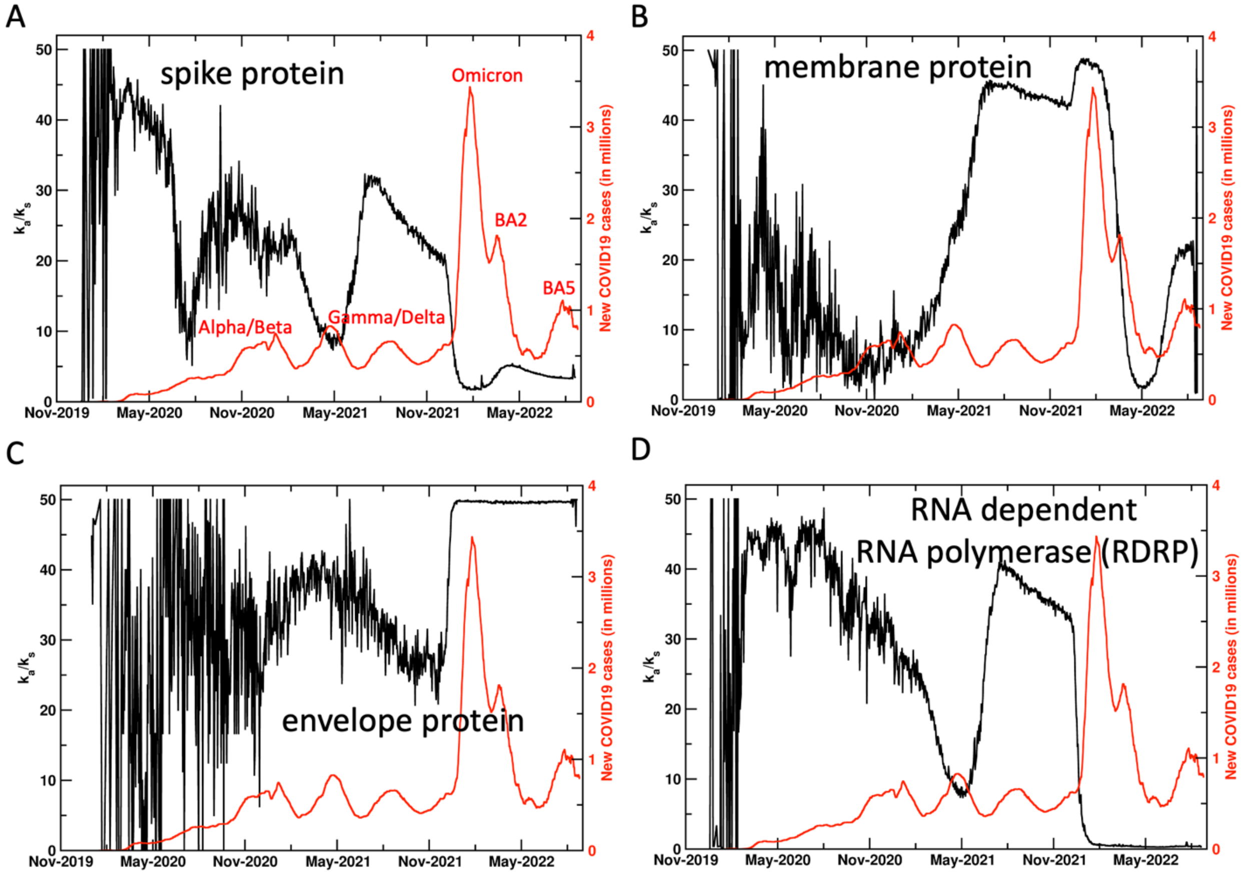


**Figure S2: Ratio of non-synonymous mutations/synonymous mutations in SARS-CoV-2 proteins.** The commonly used indicator did not provide a reliable signal for surge prediction for most proteins. The ratio for only the membrane protein shows increase before the surges associated with some variants. The information from this ratio can be used as a secondary signal to support the primary signal from $k_a$.

**Rate of mutations**

Numerical derivative of nonsynonymous mutations ($\Delta k_a/\Delta t$) calculated daily is shown in the Figure S3.

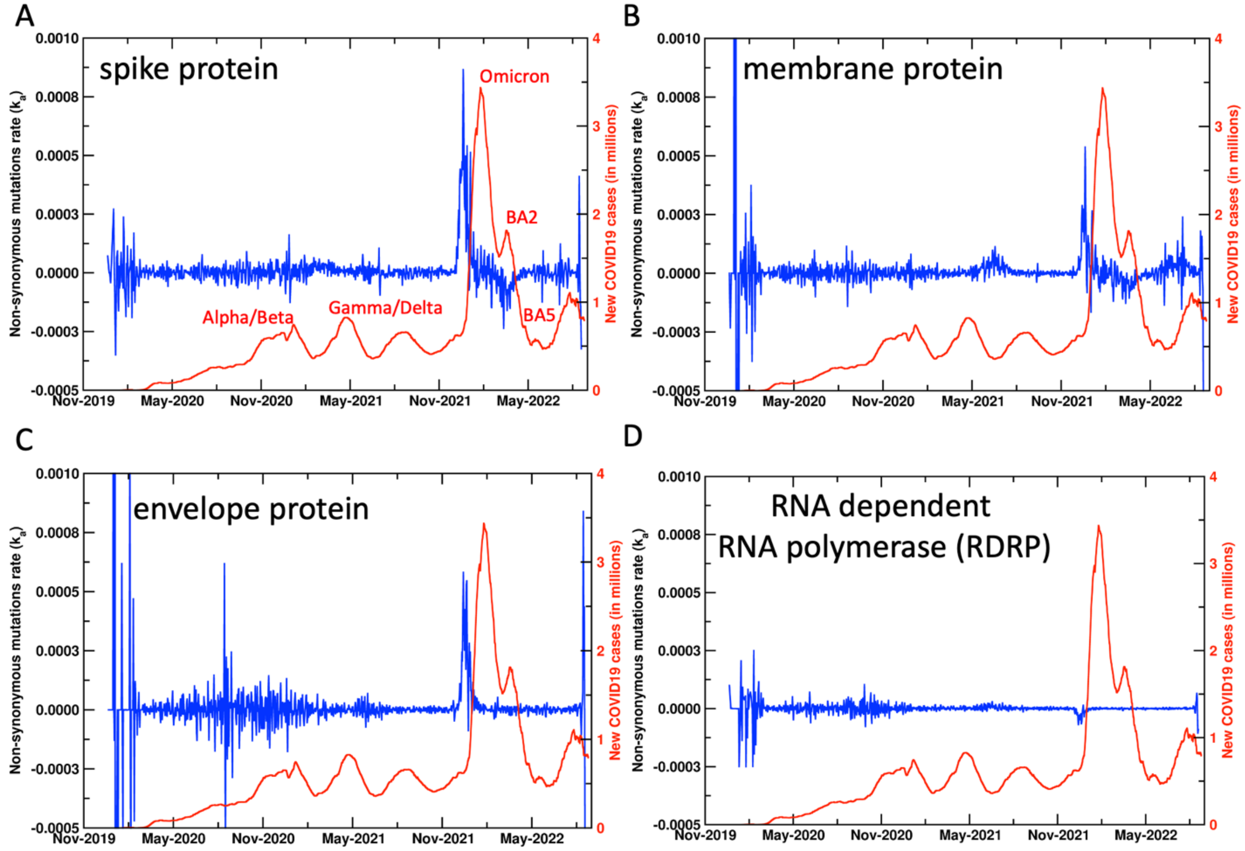


**Figure S3: Daily rate of non-synonymous mutations in SARS-CoV-2 proteins.** The rate is calculated as a numerical derivative of data shown in Figure S1. The rate shows most noticeable increase before the Omicron surge, other periods are inconclusive. Note that the nature of ongoing current data is expected to be noisy (few samples, weightings that change over days), therefore, the rate of mutations appears to be unreliable in predicting surges.
